# Olverembatinib inhibits SARS-CoV-2-Omicron variant-mediated cytokine release

**DOI:** 10.1101/2022.02.07.479443

**Authors:** Marina Chan, Eric C. Holland, Taranjit S Gujral

## Abstract

The Omicron variant has become dominant in the U.S. and around the world. This variant is found to be 2-fold more infectious than the Delta variant, posing a significant threat of severe cases and death. We and others have recently reported that the N-terminus domain (NTD) of the SARS-CoV-2 of various variants is responsible for inducing cytokine release in human PBMCs. Here, we demonstrate that the NTD of the Omicron variant remains highly effective at inducing cytokine release in human PBMCs. Furthermore, we show that Ponatinib and a novel compound, Olverembatinib, are potent Omicron NTD-mediated cytokine release inhibitors. Target profiling revealed that Olverembatinib blocks most of the previously identified kinases responsible for cytokine release. Together, we propose that Ponatinib and Olverembatinib may represent an attractive therapeutic option for treating moderate to severe COVID-19 cases.

**HIGHLIGHTS:**

- The N-terminus domain (NTD) of the SARS-CoV-2 Omicron variant strongly induces multiple inflammatory molecules in PBMCs, unaffected by the mutations observed in the NTD.
- The cytokine release mediated by the Omicron variant is comparable to the Delta variant.
- Olverembatinib, a clinical-stage multi-kinase inhibitor, potently inhibits Omicron NTD-mediated cytokine release.
- Olverembatinib could relieve severe symptoms associated with COVID-19 Omicron and Delta variants.

## Main Text

The World Health Organization has declared COVID-19 to be a pandemic. Despite the development of vaccines, COVID-19 continues to be a healthcare burden, especially in persons with a compromised immune system & others who remain unvaccinated. Most COVID-19 patients develop mild to moderate symptoms, while 15-20% of patients face hyper-inflammation induced by massive cytokine production, called ‘cytokine storm’, ultimately leading to alveolar damage and respiratory failure. Recently, we and others have shown that stimulation with S1 subunit of SARS-CoV-2 spike protein causes upregulation and release of a panel of inflammatory molecules such as IL-1b, IL-6, and CCL7 in monocytes and PBMCs (Chan et al., 2021; Kucia et al., 2021; Lu et al., 2021). Further, it has been shown that stimulation with the N-terminus domain (NTD) of the S1 subunit is sufficient to activate monocytes. Consistently, recent studies have identified several C-type lectins and Tweety family member 2 as glycan-dependent binding partners of the NTD of the SARS-CoV-2 spike protein (Lempp et al., 2021; Lu et al., 2021). The engagement of these receptors with the SARS-CoV-2 virus induces robust proinflammatory responses in myeloid cells that correlate with COVID-19 severity. Thus, identifying potential therapeutics that could abrogate NTD-mediated activation of monocytes and subsequent cytokine release is highly desirable for treating moderate to severe COVID-19 and other conditions where a cytokine storm is a lethal event.

In late November 2021, the Omicron (B. 1.1.529 / 21K) variant was detected in South Africa and has been associated with rapidly increasing case numbers worldwide (Pulliam et al., 2021). The Omicron variant carries more than 50 mutations, including more than 25 mutations in the Spike protein alone. Of these, ~30% of mutations are present in the NTD. Here, we sought to determine whether the mutations in the NTD of the Omicron variant affect its ability to activate myeloid cells and promote the secretion of inflammatory cytokines. First, we stimulated pooled peripheral blood mononuclear cells (PBMCs) with mammalian cell (HEK293)-derived NTD (1μg/mL) of the Omicron variant. We observed a robust (>100-fold) increase in the release of interleukins including IL-1β, IL-6, and tumor necrosis factor (TNFα) (**Fig. 1B**), suggesting that the Omicron variant NTD can still activate myeloid cells and promote cytokine release. Next, we compared the changes in cytokine release in PBMCs produced by the stimulation of NTD from the Omicron, Delta, and Wuhan variants. Our data show that the NTD from the Omicron variant was equally effective in promoting cytokine release. Together, these data establish that the presence of mutations in the NTD of the Omicron variant does not alter its ability to promote cytokine release.

**Figure 1.**
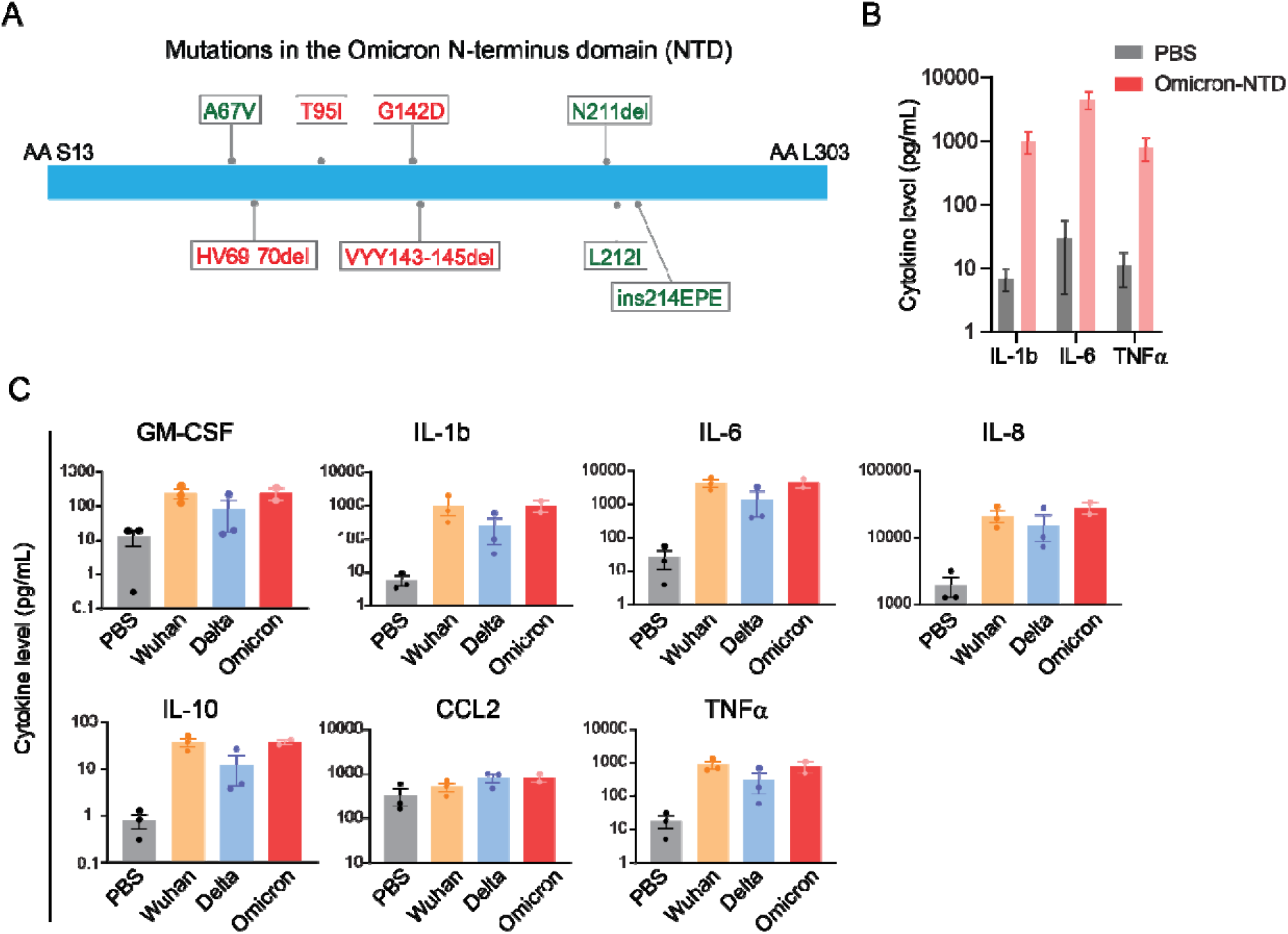
NTD of the SARS-CoV-2 Omicron variant stimulates cytokine release. ***A.*** A schematic showing mutation in the Omicron NTD. Unique mutations found in the Omicron variant are shown in green color. ***B.*** Comparison of changes in cytokine release in response to the recombinant NTD from Wuhan, Delta, and Omicron variants. Measurement of cytokine release from healthy donor PBMCs treated with PBS or Omicron NTD at 1 μg/ml for 24 h. ***C.*** Comparison of cytokine release from healthy donor PBMCs treated with Wuhan, Delta, or Omicron NTD at 1 μg/ml for 24 h. Cytokine release in the conditioned media was measured by Luminex. Recombinant NTD of different variants were purified from HEK293 cells. Data are shown as the mean of two to three biological replicates. Error bars denote SEM.

Previously, a machine learning-based drug screening identified Ponatinib, an FDA-approved drug for chronic myelogenous leukemia, as a potent inhibitor of S1 protein-mediated cytokine release in PBMCs. (Chan et al., 2021). Thus, we asked whether Ponatinib treatment could also inhibit cytokine release mediated by the NTD from the Omicron variant. In addition, we also evaluated the efficacy of Baricitinib, an FDA-approved JAK inhibitor for the treatment of COVID-19 (Favalli et al., 2020), and Olverembatinib, a clinical-stage multi-kinase inhibitor, that is structurally similar to ponatinib, with a manageable safety profile (Jiang et al., 2019). Our data show that treatment with Ponatinib or Olverembatinib inhibited the NTD-mediated release of all seven cytokines measured in a dose-dependent manner (**Fig. 2A**). Both Ponatinib and Olverembatinib inhibited the release of cytokines even at low nanomolar concentrations (< 50 nM). In contrast, treatment with Baricitinib only inhibited three out of seven cytokines measured (GM-CSF, CCL2, and IL-10) at 1000 nM (**Fig. 2A**). Since Olverembatinib treatment showed the most substantial suppression of the Omicron-NTD-mediated cytokine release, we next determined the Olverembatinib concentration range for the panel of cytokines tested, required to inhibit 50% of the Omicron NTD-mediated cytokine release (EC_50_), was between 7.7 and 56 nM (**Fig. 2B**). Previously, several protein kinases, including JAK1, EPHA7, IRAK1, MAPK12, and MAP3K3, were identified as essential for S1 protein-mediated cytokine release in myeloid cells (Chan et al., 2021). The kinase activity profile of Olverembatinib showed that this drug inhibits 11 out of 13 kinases predicted to be essential for the NTD-mediated chemokine and cytokine release. Thus, these data indicate that Olverembatinib, a clinical-grade, multi-specific kinase inhibitor, blocks the activity of several kinases essential for cytokine signaling, thereby dampening the Omicron NTD-mediated cytokine release.

**Figure 2.**
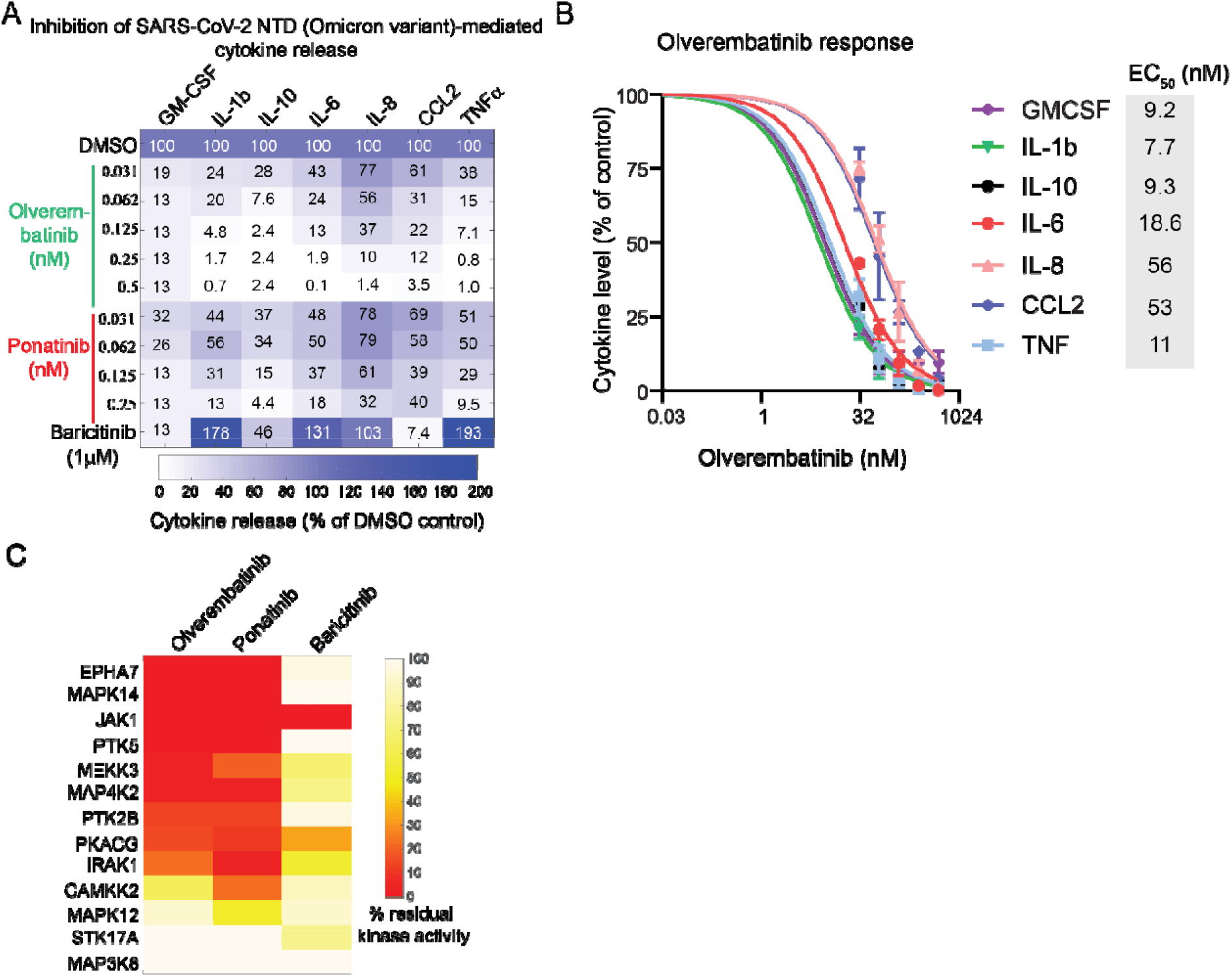
Olverembatinib is a potent inhibitor of Omicron NTD-mediated cytokine release. ***A.*** Effect of Olverembatinib, Ponatinib, and Baricitinib on Omicron NTD-mediated cytokine release. ***B.*** EC_50_ of Olverembatinib on indicated cytokines. ***C.*** Comparison of kinase inhibition profiles of Olverembatinib, Ponatinib, and Baricitinib. PBMCs from healthy donors were treated with Omicron NTD at 1 μg/ml and Olverembatinib, Ponatinib, or Baricitinib at indicated concentrations. Cytokine release in the conditioned media was measured by Luminex. Data are shown as the mean of two biological replicates. Error bars denote SEM.

Taken together, we propose that agents targeting multiple kinases required for SARS-CoV-2-mediated cytokine release, such as Olverembatinib and Ponatinib, may represent an attractive therapeutic option for treating moderate to severe COVID-19.

## Notes

### Competing Interest Statement

The authors have declared no competing interest.

